# Optimization of DNA Transformation in *Mesoplasma florum* and Identification of a Candidate Recipient Strain for Genome Transplantation

**DOI:** 10.64898/2026.09.29.755066

**Authors:** Jeremy Gagnon, Dominick Matteau, Frederic Grenier, Sébastien Rodrigue

## Abstract

Genome transplantation is a key technology for synthetic genomics, enabling entire genomes to be transferred into recipient cells. Despite its importance, genome transplantation remains confined to a small number of wall-less Mollicute species and is often characterized by low efficiencies, limiting the development and testing of synthetic genomes. Because genome transplantation relies on the same polyethylene glycol mediated DNA delivery process used for plasmid transformation, improving DNA uptake is an important step toward more efficient transplantation systems. Here, we systematically optimized polyethylene glycol-mediated transformation in *Mesoplasma florum*, a fast-growing, non-pathogenic Mollicute with a reduced genome that represents an attractive chassis for synthetic genomics. We evaluated 20 parameters spanning cell physiology, DNA preparation, membrane conditioning, and recovery conditions. DNA topology emerged as the strongest determinant of transformation efficiency, with highly compacted DNA preparations producing up to three orders of magnitude more transformants than conventional plasmid preparations. Growth phase and polyethylene glycol concentration also strongly influenced transformation outcomes. To identify potential recipients for future genome transplantation experiments, we further screened ten strains belonging to the *Mesoplasma* lineage. Transformation efficiencies varied widely among strains, and *Mesoplasma entomophilum* W17 emerged as a particularly promising candidate. Moreover, W17 supported replication of plasmids carrying the *M. florum oriC*, suggesting compatibility between the replication systems of the two species. Together, these results establish an improved transformation workflow for *M. florum*, identify DNA topology as a major determinant of DNA uptake, and reveal a new candidate recipient for future genome transplantation studies. These advances provide practical tools and biological insights for synthetic genome engineering in Mollicutes.

## Introduction

Genome transplantation enables a complete genome to be transferred from a donor cell into a recipient cell, thereby replacing the genetic information that controls cellular functions. This technology underpinned the construction of JCVI-syn1.0, the first cell controlled by a chemically synthesized genome derived from *Mycoplasma mycoides*, and the subsequent development of the minimal cell JCVI-syn3.0, which retains the same genomic lineage.^1–3^ Beyond demonstrating the feasibility of large-scale genome design and synthesis, genome transplantation provides a practical route for constructing and evaluating extensively engineered chromosomes that cannot readily be assembled or tested using conventional genetic approaches. As synthetic genomics continues to expand, genome transplantation remains one of the few technologies capable of converting a designed genome into a living cell.

Most successful genome transplantation experiments have been performed in members of the class Mollicutes, a group of bacteria characterized by the absence of a cell wall and relatively small genomes (0.5 to 2.2 Mb).^4^ The wall-less nature of these organisms facilitates polyethylene glycol (PEG)-mediated membrane fusion, a process that underlies both transformation and genome transplantation. Combined with their reduced cellular complexity, these features have established Mollicutes as the principal experimental platform for synthetic genome engineering and minimal-cell research. Despite these advantages, genome transplantation has only been demonstrated in a limited number of donor-recipient species combinations, and the factors governing recipient permissiveness remain poorly understood.^5,6^

Among Mollicutes, *Mesoplasma florum* has emerged as an attractive candidate for synthetic genomics. Unlike several species traditionally used in transplantation studies, *M. florum* is not associated with human, livestock, or plant diseases. *M. florum* combines a relatively small genome of approximately 0.79 Mb with rapid growth under laboratory conditions, exhibiting a doubling time of approximately 32 minutes.^7^ Over the last decade, the development of genetic tools, genome engineering methods, essentiality datasets, and systems-level genomic resources has substantially expanded our ability to manipulate and model this organism.^8–12^ Together, these advances have strengthened its potential as a chassis for synthetic genome design, genome reduction, and the study of fundamental cellular processes.

Despite its promise, efficient introduction of exogenous DNA remains a persistent challenge in *M. florum* and in Mollicutes more broadly. PEG-mediated transformation remains the most widely used method for introducing DNA into these organisms, yet reported transformation efficiencies are often highly variable both within and between species.^13^ The mechanism is thought to involve dehydration-induced membrane fusion, during which DNA becomes associated with the cell surface and is subsequently internalized upon rehydration.^14,15^ Because genome transplantation relies on the same PEG-mediated DNA delivery process, limitations affecting plasmid transformation are likely to become even more restrictive when entire chromosomes are used. Understanding the parameters that govern DNA uptake therefore represents an important step toward improving both routine genetic manipulation and future genome transplantation workflows.

In addition to transformation efficiency itself, the choice of recipient species may strongly influence transplantation success. Previous transplantation of the *M. florum* genome into *Mycoplasma capricolum* demonstrated the feasibility of generating cells controlled by an *M. florum* genome, but the efficiency of this process remained low, reaching only approximately 3 transplants µg⁻¹ of carefully prepared *M. florum* genomic DNA and 21 transplants µg⁻¹ of a purified *M. florum* genome cloned and maintained in yeast.^9^ One possible explanation is the substantial evolutionary distance separating these species. If recipient permissiveness is influenced by phylogenetic relatedness, species within the *Mesoplasma* lineage may provide more compatible recipients for *M. florum* genome transplantation than those identified to date. Expanding the range of potential recipients could therefore facilitate future efforts to engineer synthetic *Mesoplasma* genomes and improve our understanding of the factors governing successful genome transfer.

In this study, we addressed two limitations that currently constrain genome engineering in *M. florum*: inefficient DNA delivery and the lack of candidate recipient species closely related to *M. florum*. We systematically optimized PEG-mediated transformation and evaluated additional *Mesoplasma* species as prospective recipients for future genome transplantation experiments. Together, these efforts aimed to establish a stronger experimental foundation for synthetic genome engineering within this group of wall-less bacteria.

## Results and Discussion

### Plasmid systems for evaluating DNA delivery and maintenance in *M. florum*

To investigate factors limiting genetic manipulation in *M. florum*, we first established a set of genetic tools that enabled successful DNA delivery to be evaluated across distinct maintenance mechanisms. Because transformation efficiency reflects not only DNA entry into the cell but also the subsequent fate of the introduced molecule, comparing plasmids with different modes of maintenance was important for interpreting the effects of protocol optimization. We therefore employed three complementary plasmid systems differing in their mode of maintenance within the recipient cell (Fig. 1). The first relies on autonomous replication through the native chromosomal origin, *oriC*. Plasmid pMflT-o4 carries the *M. florum oriC* region and can be maintained as an extrachromosomal replicon following transformation, although homologous recombination between the plasmid and chromosomal *oriC* regions can also generate integrated forms (Fig. 1A).^8^ The second system uses Tn4001-mediated transposition.^16–19^ Plasmids pDM017 to pDM023 encode the transposition machinery required for random chromosomal insertion of a selectable marker. Following evaluation of several promoter and antibiotic-resistance gene combinations (Table S1), pDM023 yielded the highest transformation efficiencies and was selected for subsequent experiments (Fig. 1B). The third system enables site-specific chromosomal integration through the Bxb1 serine integrase. Plasmid pJG001 carries a Bxb1 *attP* site and integrates into a pre-positioned chromosomal *attB* landing site in previously described strain MFSC001^20^, enabling targeted insertion at a defined genomic locus (Fig. 1C and Fig. S1).^21,22^ In addition, pJG001 carries the *E. coli oriC*, allowing the plasmid to be processed using the OriCiro system and thereby providing an opportunity to investigate the influence of DNA topology on transformation efficiency.^23,24^

**Figure 1:**
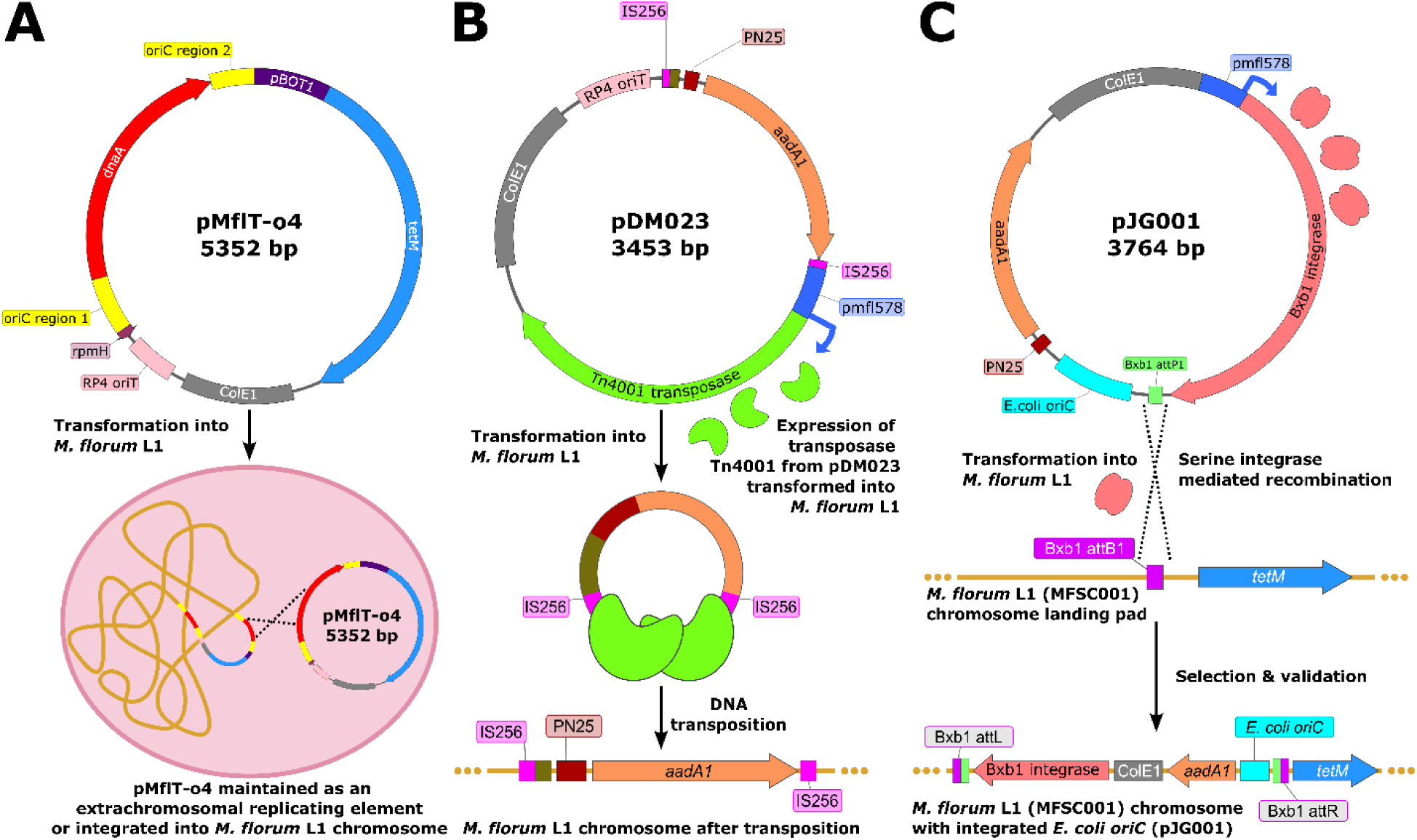
Plasmid systems used in this study **A)** pMfIT-o4 plasmid containing *M. florum oriC*. Upon transformation into *M. florum* L1, pMfIT-o4 is maintained either as an extrachromosomal replicating element or through chromosomal integration. Adapted from Matteau *et al*.^8^ **B)** pDM023 plasmid carrying an engineered copy of the Tn4001 transposase coding gene. After transformation into *M. florum* L1, expression of the Tn4001 transposase drives transposition of a selectable cassette into the host chromosome, generating stable integrants. **C)** pJG001 plasmid, which includes an *E. coli oriC*, a Bxb1 serine integrase and its attP1 site, transformed into *M. florum* L1 MFSC001, and leading to site-specific recombination at the chromosomal Bxb1 attB1 landing pad.

These systems provided complementary readouts of DNA delivery and maintenance. For transformation optimization, their combined use enabled parameters affecting successful DNA introduction to be evaluated across distinct genetic contexts. For strain-screening experiments, the transposition-based system offered a broadly applicable transformation readout across diverse *Mesoplasma* backgrounds, whereas the *oriC*-based constructs simultaneously reported on replication compatibility between species, a property directly relevant to genome transplantation. Together, these genetic tools established the experimental framework used throughout this study.

### Optimization of PEG-mediated transformation in *M. florum*

To evaluate DNA delivery in *M. florum*, we employed a PEG-mediated transformation workflow adapted from previously described protocols (Fig. 2).^8^ Cells were conditioned with CaCl₂^25,26^, exposed to plasmid DNA, treated with PEG-containing fusion buffer, and allowed to recover before selection. The parameters evaluated in our optimization campaign were selected to test hypotheses concerning the recipient cell, the transformation environment, and the incoming DNA. We systematically examined 20 parameters spanning these different aspects of the workflow (Table 1 and Table S1).

**Figure 2:**
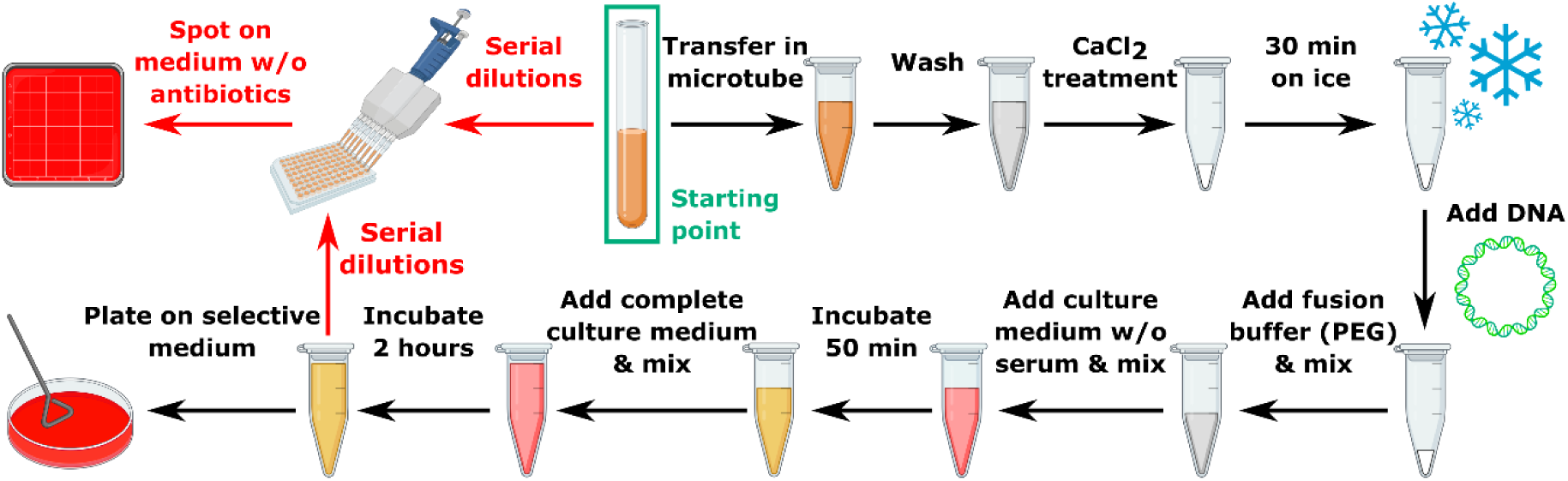
Workflow for PEG-mediated transformation The PEG transformation workflow consists of sequential steps beginning with cell harvesting and washing, followed by CaCl₂ treatment and incubation on ice to promote competence. Plasmid DNA is then added, and fusion is induced by mixing with PEG-based fusion buffer. After addition of serum-free medium and a 50-min incubation, complete medium is supplied and cells recover for 2 h at the appropriate temperature. The culture is subsequently diluted and spotted on non-selective medium before plating on antibiotic-containing agar to isolate transformants. The initial culture is diluted and spotted on a non-selective medium to measure cell concentrations at the beginning of the workflow.

**Table 1:**
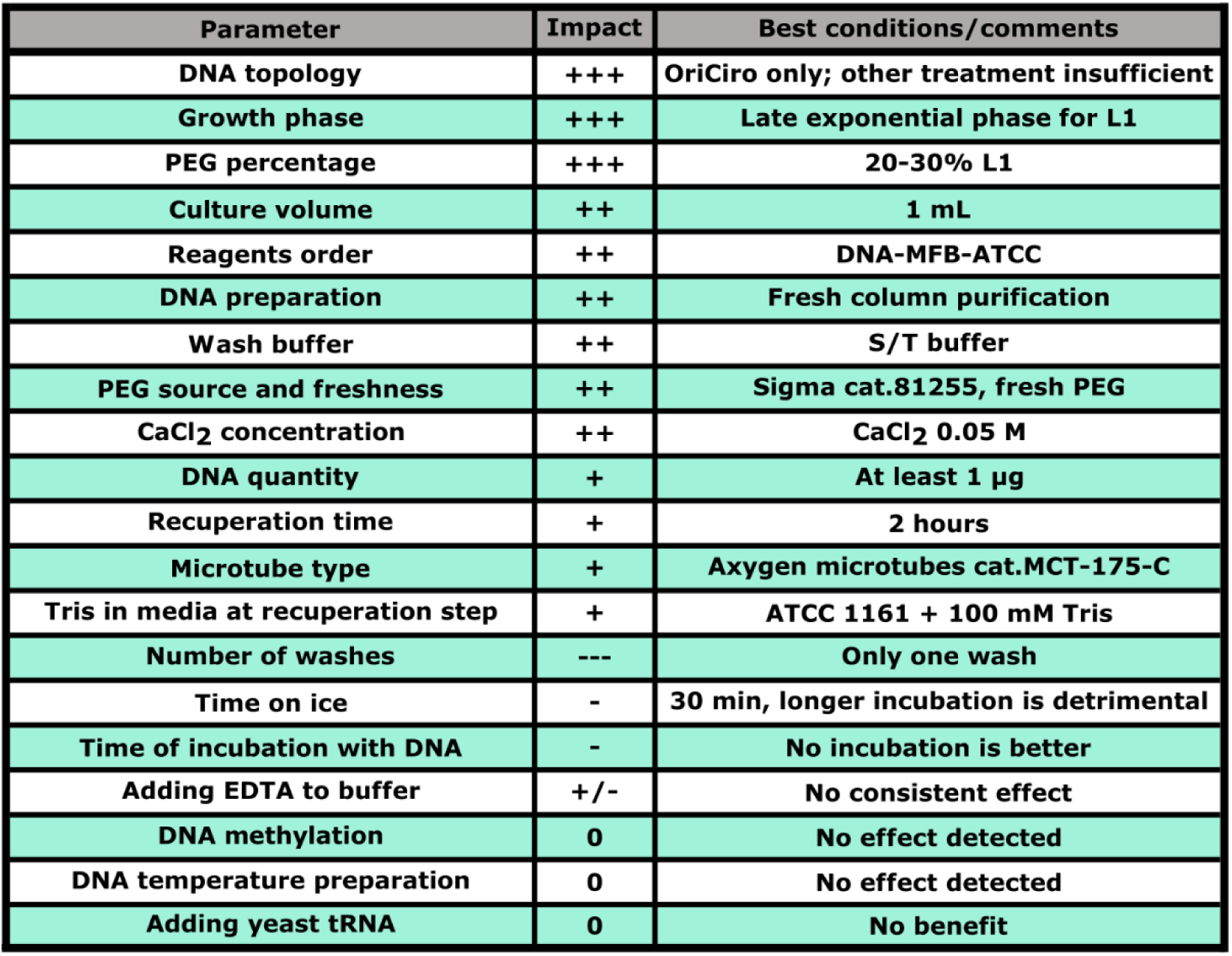
Key factors influencing PEG-mediated transformation. Summary of the main experimental parameters that affect PEG-mediated transformation efficiency, highlighting the dominant influence of DNA topology, strain background, PEG quality, and preparation conditions.

### Highly supercoiled OriCiro-amplified DNA markedly enhances transformation efficiency

Among the properties of the incoming DNA, we hypothesized that highly supercoiled molecules would be delivered more efficiently during PEG-mediated transformation. Supercoiled DNA adopts a more compact conformation than relaxed circular DNA, thereby reducing its effective dimensions (Fig. 3A).^27,28^ We reasoned that this compaction could facilitate DNA transfer across the membrane of recipient cells. To test this hypothesis, we used pJG001 (Fig. 1C), which contains the *E. coli oriC* sequence required for amplification using the OriCiro cell-free replication system. OriCiro reconstitutes the *E. coli* chromosomal replication cycle in vitro using 26 purified replication proteins, enabling exponential amplification of circular DNA carrying *oriC* and generating highly supercoiled plasmid products (Fig. S2)^23,24^. We therefore compared the transformation efficiency of OriCiro-amplified pJG001 with that of pJG001 extracted directly from *E. coli*.

**Figure 3:**
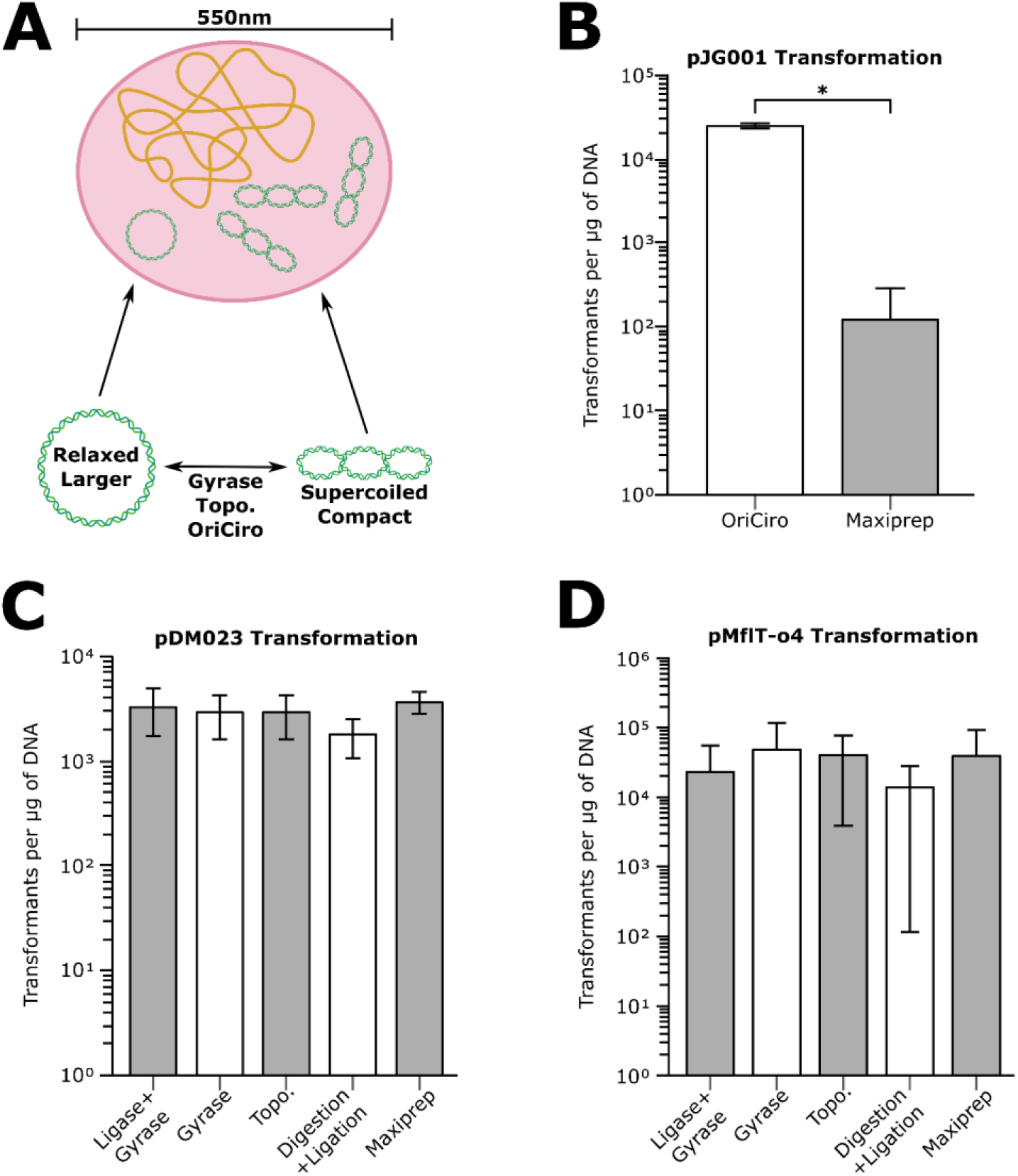
Impact of DNA topology and preparation methods on transformation efficiency in *Mesoplasma florum* L1. **A)** Schematic representation of relaxed versus supercoiled DNA topologies. Gyrase and topoisomerase modulate DNA compaction, generating either extended relaxed molecules or highly supercoiled compact forms. **B)** Transformation efficiency of plasmid pJG001 prepared using OriCiro versus untreated Maxiprep. OriCiro-treated DNA yielded significantly higher transformation levels (p < 0.05). **C)** Transformation of pDM023 following different DNA preparation workflows. **D)** Transformation of pMfIT-o4 across the same preparation conditions.

OriCiro-amplified pJG001 produced nearly three orders of magnitude more transformants in *M. florum* than conventionally prepared pJG001 (Fig. 3B). This difference was statistically significant (Welch’s two-sample *t*-test, *p* = 0.036) and represented the largest effect observed among the 20 parameters evaluated. These results support our hypothesis that the physical conformation of incoming DNA strongly influences successful DNA delivery during PEG-mediated transformation.

Because the OriCiro kits became commercially unavailable following the acquisition of OriCiro Genomics by Moderna in 2023, we investigated whether the effect could be reproduced using commercially available enzymes that modify plasmid topology.^24^ Plasmid preparations were treated with *E. coli* DNA gyrase, either alone or following ligation, whereas topoisomerase I was used to generate more relaxed DNA. We also produced relaxed circular DNA by restriction digestion followed by ligation and compared these preparations with untreated maxiprep DNA. None of these treatments reproduced the large increase in transformation efficiency observed with OriCiro-amplified pJG001 (Fig. 3B-D; Table S1). Moreover, transformation efficiencies did not differ significantly among treatments for pDM023 or pMflT-o4 (one-way ANOVA, *p* = 0.637 and *p* = 0.963, respectively).

The inability of gyrase treatment to reproduce the OriCiro effect indicates that post-purification treatment with this enzyme was insufficient under the conditions tested. This result does not contradict the proposed role of DNA compaction because OriCiro generates plasmid DNA through a complete replication cycle, whereas gyrase treatment modifies DNA that has already been replicated and purified from *E. coli*. The two processes may therefore produce different distributions of topological forms. Consistent with a more complex relationship between enzymatic treatment and transformation efficiency, topoisomerase I occasionally produced transformation efficiencies comparable to those obtained with gyrase despite promoting DNA relaxation. The highly supercoiled DNA generated by OriCiro amplification was therefore particularly favorable for PEG-mediated transformation, although this phenotype could not be reproduced by treating conventionally prepared plasmids with individual topology-modifying enzymes.

### Growth phase and PEG concentration define optimal competence and transformation conditions

Having examined the physical properties of the incoming DNA, we next evaluated factors associated with the recipient cells and the transformation environment. The physiological state of the culture had a pronounced effect on transformation efficiency. Cultures harvested during late exponential phase yielded 3.8 × 10⁴ transformants µg⁻¹ DNA, compared to 1.4 × 10³ and 8.8 × 10³ transformants µg⁻¹ DNA for cultures harvested during early and mid-exponential phase, respectively (Table 1 and Table S1). Late-exponential cultures therefore produced approximately 27-fold and fourfold more transformants than early- and mid-exponential cultures, respectively. The highest efficiencies were obtained within a narrow window of culture coloration, from the onset of yellowing to a fully yellow medium, corresponding to mid- to late exponential phase (Fig. S3). This result contrasts with the optimum previously reported for *M. capricolum*, in which early exponential phase is favored for both plasmid transformation and genome transplantation.^13,29^ The physiological window associated with competence thus appears to differ among Mollicute species, emphasizing the need to re-establish this parameter when adapting transformation protocols to a new recipient.

PEG concentration was the strongest determinant associated with the transformation environment. Because PEG promotes the membrane dehydration and fusion required for DNA transfer, we tested concentrations ranging from 10 to 40% (wt/vol). Transformation efficiency showed a narrow optimum at 20% PEG, reaching 7.6 × 10⁴ transformants µg⁻¹ DNA (Table 1 and Table S1). Efficiency decreased to 2.7 × 10⁴ transformants µg⁻¹ DNA at 15% PEG, 1.1 × 10⁴ transformants µg⁻¹ DNA at 25% PEG, and 5.5 × 10³ transformants µg⁻¹ DNA at 40% PEG, corresponding to approximately threefold, sevenfold, and fourteenfold reductions relative to the optimum, respectively. PEG source also affected transformation, with the MilliporeSigma PEG 6000 (Sigma 81255) product yielding approximately fourfold more transformants than the PEG 8000 (Biolynx UB19959S2) product under otherwise comparable conditions. These results indicate that PEG-mediated transformation requires tightly controlled conditions and that PEG concentration and source should be treated as critical experimental variables rather than interchangeable protocol details. The sharp optimum is consistent with a balance between insufficient membrane perturbation at lower PEG concentrations and increasingly unfavorable effects on recipient cells at higher concentrations.

### Membrane conditioning and the timing of DNA-cell contact influence transformation efficiency

We next examined parameters affecting the condition of the cell surface and its interaction with DNA. The order in which reagents were added after CaCl₂ treatment had a pronounced effect on transformation efficiency. Using pMflT-o4 under otherwise comparable conditions, adding DNA directly to the calcium-conditioned cells before the PEG-containing fusion buffer yielded 8.8 × 10⁵ transformants µg⁻¹ DNA, nearly fivefold more than when DNA was premixed with the fusion buffer before addition to the cells, which yielded 1.8 × 10⁵ transformants µg⁻¹ DNA. An alternative order of addition produced an intermediate efficiency of 3.0 × 10⁵ transformants µg⁻¹ DNA (Table 1 and Table S1). These results suggest that contact between DNA and the calcium-conditioned cell surface before PEG treatment favors efficient transformation, whereas exposing DNA to the PEG-containing buffer before its addition to the cells is less favorable.

Wash-buffer composition also influenced transformation efficiency. We compared S/T buffer, phosphate-buffered saline, and a HEPES-based electroporation buffer, each with or without 10 mM EDTA. S/T buffer produced the highest transformation efficiency, reaching 1.3 × 10⁵ transformants µg⁻¹ DNA, compared with 7.0 × 10⁴ transformants µg⁻¹ DNA for phosphate-buffered saline and 3.0 × 10⁴ transformants µg⁻¹ DNA for the electroporation buffer (Table S1). EDTA supplementation produced no consistent improvement across the conditions tested. EDTA had been included to chelate divalent cations and potentially limit nuclease activity, but the absence of a measurable benefit indicates that EDTA-mediated nuclease inhibition was not beneficial under these conditions.

CaCl₂ concentration and incubation time required similar control. Among the concentrations tested, 50 mM CaCl₂ produced the highest and most reproducible transformation efficiency, yielding 4.5 ± 1.6 × 10² transformants µg⁻¹ DNA across replicate experiments, compared with 1.2 ± 0.4 × 10² and 1.5 ± 0.6 × 10² transformants µg⁻¹ DNA at 10 and 100 mM CaCl₂, respectively (Table 1 and Table S1). Extending the incubation with CaCl₂ from 30 to 60 min reduced transformation efficiency from 4.5 × 10² to 2.0 × 10² transformants µg⁻¹ DNA. Moreover, introducing a second wash after CaCl₂ treatment reduced transformation efficiency to near the detection limit. This pronounced decrease is consistent with disruption of the calcium-conditioned state established before DNA addition.

The interval between DNA addition and PEG treatment was also examined. A 5- or 15-min incubation of cells with DNA before addition of the fusion buffer yielded slightly lower transformation efficiencies of 8 and 10 transformants µg⁻¹ DNA, respectively, compared with 16 transformants µg⁻¹ DNA when the fusion buffer was added immediately after DNA (Table S1). Although this modest difference should be interpreted cautiously given the small numbers of transformants, the results provided no evidence that extending the interval between DNA addition and PEG treatment improves transformation.

These findings demonstrate that membrane conditioning and the order and timing of reagent addition are important determinants of PEG-mediated transformation. Transformation efficiency depends not only on reagent composition but also on the sequence and timing with which cells are exposed to DNA and PEG.

### Culture volume, DNA input quantity, and plasmid preparation method influence transformation efficiency

We next examined how recipient culture volume and DNA input quantity affected transformation efficiency. Cultures of 0.2, 1, and 10 mL were tested with different amounts of plasmid DNA (Table 1 and Table S1). Within each experimental series, using 1 mL of culture produced the highest transformation efficiency. Reducing the culture volume to 0.2 mL decreased transformation efficiency by approximately five- to ninefold, from 1.3 × 10³ to 1.7 × 10³ transformants µg⁻¹ DNA with 1 mL of culture to 1.4 × 10² to 3.5 × 10² transformants µg⁻¹ DNA with 0.2 mL. Conversely, increasing the culture volume from 1 to 10 mL reduced transformation efficiency from 7.6 × 10⁴ to 1.9 × 10⁴ transformants µg⁻¹ DNA. Increasing culture volume therefore did not proportionally improve transformation, and an intermediate culture input produced the highest efficiency under the reaction conditions tested.

Increasing the DNA input quantity from 1 to 10 µg increased the absolute number of transformants but reduced transformation efficiency per unit of DNA from 1.3 × 10³ to 5.9 × 10² transformants µg⁻¹ DNA (Table 1 and Table S1). Higher DNA inputs therefore produced diminishing returns rather than proportional increases in transformation.

The plasmid preparation method also influenced transformation outcomes. Using pDM023, DNA obtained by either classic home-made chloroform purification or commercial column miniprep procedures produced few or no transformants, with efficiencies ranging from 0 to 2 transformants µg⁻¹ DNA. In the same experimental series, maxiprep preparations yielded 80 to 2.6 × 10² transformants µg⁻¹ DNA (Table 1 and Table S1). Although the absolute yields in this series were lower than those obtained in some of the other optimization experiments, the consistent within-experiment difference between miniprep and maxiprep DNA supported the use of maxiprep preparations for subsequent transformations. Neither the age nor heat treatment of maxiprep DNA produced an obvious loss of performance. Performing DNA preparation at 4 °C, which was tested as a possible means of preserving supercoiled plasmid forms, also produced no detectable improvement. Maxiprep DNA therefore provided a substantially more reliable substrate than miniprep DNA for PEG-mediated transformation. Its improved performance likely results from more effective removal of residual salts and cellular contaminants that could alter the ionic environment of the transformation reaction or interfere with interactions between plasmid DNA and the calcium-conditioned cell surface.

These experiments revealed diminishing returns from increasing either culture volume or DNA input quantity and identified plasmid preparation method as an important determinant of transformation performance.

### Recovery conditions and secondary protocol variables

Post-transformation recovery conditions were examined to determine how long cells should be incubated before selection. Recovery for 2 h produced a higher transformation efficiency than recovery for either 1 or 3 h, yielding 4.5 × 10² transformants µg⁻¹ DNA compared with 2.8 × 10² and 3.0 × 10² transformants µg⁻¹ DNA, respectively (Table 1 and Table S1). Supplementing the recovery medium with 100 mM Tris further increased transformation efficiency to 5.0 × 10² transformants µg⁻¹ DNA. These results are consistent with a balance between allowing sufficient time for transformed cells to recover and express the selectable marker while limiting the effects of prolonged incubation in an increasingly acidified medium. The modest improvement associated with Tris supplementation is also consistent with a beneficial effect of increased buffering capacity.

Several additional protocol variables produced modest or inconsistent effects. Axygen microcentrifuge tubes (cat. MCT-175-C) yielded 8.4 × 10² to 9.4 × 10² transformants µg⁻¹ DNA, compared with 5.0 × 10² to 6.8 × 10² transformants µg⁻¹ DNA using VWR microtubes (cat. 87003- 294) (Table S1). This difference may reflect variation in cell or DNA adsorption to the tube surface during centrifugation and resuspension. Vortex mixing also produced a slightly higher transformation efficiency than gentle inversion, with 68 compared with 40 transformants µg⁻¹ DNA. However, these relatively small differences should be interpreted as practical refinements rather than major determinants of transformation efficiency. Transferring the protocol from 50-mL tubes to a microcentrifuge-tube format did not appreciably alter performance but simplified handling and reduced reagent use.

Other modifications provided no detectable benefit. Plasmid pDM023 prepared in either DH5α *pir*+ or the methylase-deficient strain ER2925 produced similarly low transformation efficiencies, indicating that removal of Dam and Dcm methylation did not improve transformation under the conditions tested. Supplementation with yeast tRNA as a potential DNA carrier also failed to improve transformation.

These results distinguish recovery duration as a meaningful component of the transformation workflow, whereas most secondary modifications produced modest, inconsistent, or undetectable effects.

### Operator-dependent execution contributes to experimental variability

Substantial variability persisted among experiments performed under nominally identical conditions, with transformation efficiencies differing by more than one order of magnitude in some comparisons (Table 1 and Table S1). Transformation efficiencies also differed substantially among experiments performed by different operators under nominally similar conditions. This observation indicates that standardizing reagent composition and incubation conditions alone may be insufficient to ensure reproducible PEG-mediated transformation.

Several steps in the protocol involve operator-dependent manipulations that are difficult to standardize completely. In particular, *M. florum* cells form aggregates that must be disrupted during washing and resuspension, and differences in aggregate dispersion may affect both cell recovery and exposure to DNA. The efficiency of pellet resuspension and the order and timing of DNA and fusion-buffer addition may also vary among operators. These differences in execution provide plausible explanations for the observed variability.

The optimized protocol therefore specifies not only reagent concentrations and incubation times but also the execution of the most sensitive handling steps. Thorough and consistent resuspension of cell pellets, complete disruption of visible aggregates, and rapid progression from DNA addition to PEG treatment are particularly important for reproducibility. The microcentrifuge-tube format also simplified handling and reduced the number of transfers required during the procedure.

Operator-dependent variability may contribute to the broad range of transformation efficiencies reported for Mollicutes across experiments and laboratories. Detailed protocol documentation and consistent execution of handling-sensitive steps are therefore particularly important for genome transplantation, where low baseline efficiencies increase sensitivity to variation in DNA delivery.

### Recommended conditions for PEG-mediated transformation of *M. florum*

The optimization campaign defined a set of recommended conditions for PEG-mediated transformation of *M. florum*. Cultures should be harvested during late exponential phase, corresponding to a translucent yellow color (Fig. S3), and 1 mL of culture should be washed once with S/T buffer before incubation in 50 mM CaCl₂ for 30 min. One microgram of maxiprep-quality plasmid DNA should then be added directly to the conditioned cells, followed immediately by fusion buffer containing 20% PEG. Cells should be allowed to recover for 2 h before selection. Cell pellets should be thoroughly resuspended, visible aggregates should be disrupted, and the interval between DNA and PEG addition should be minimized.

Among the variables examined, OriCiro amplification produced the largest increase in transformation efficiency, although the commercial system is no longer available. For conventionally prepared plasmids, maxiprep DNA provided the most reliable substrate. PEG concentration and source, culture growth phase, membrane conditioning, and reagent-addition sequence should be controlled carefully. By contrast, EDTA supplementation, altered DNA methylation, yeast tRNA supplementation, and low-temperature DNA preparation provided no measurable benefit. Tube selection, mixing method, Tris supplementation during recovery, and other secondary variables produced comparatively modest effects.

These recommendations provide a standardized framework for PEG-mediated transformation in *M. florum* while recognizing that operator-dependent execution remains an important source of variability. Because several of the strongest effects involved properties of the recipient cells, we next asked whether transformation competence varies among related strains within the *Mesoplasma* lineage.

### Transformation efficiency varies widely across *Mesoplasma* strains

To expand the range of potential recipients for future transplantation of the *M. florum* L1 genome, we next evaluated whether related *Mesoplasma* strains could be transformed efficiently. Using the optimized PEG-mediated transformation protocol, we screened ten strains, including *M. florum* L1, *M. entomophilum* W17, and two isolates of *Mesoplasma coleopterae*, BARC 781 and BARC 786 (Fig. 4A). Each strain was transformed with pDM023, which integrates a selectable marker into the chromosome through Tn4001-mediated transposition. Although this readout depends on both DNA delivery and transposition, it avoids the requirement for a host-compatible origin of replication and therefore provides a common basis for comparing transformation across strains.

**Figure 4:**
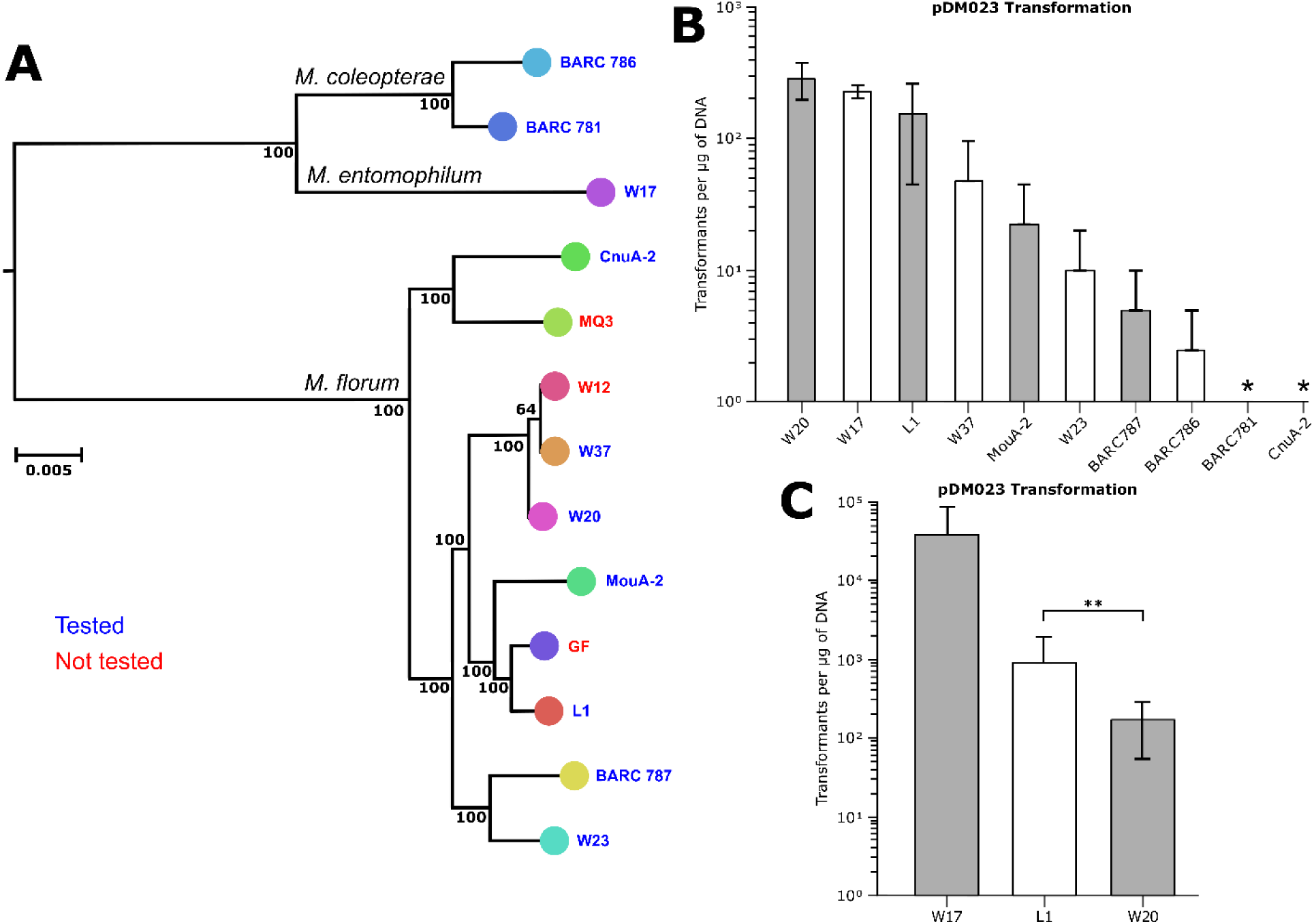
Transformation efficiency across ten *Mesoplasma* strains. **B)** *Mesoplasma* phylogenetic tree constructed using concatenated alignments of 412 conserved proteins.^11^**B)** PEG-mediated transformation efficiency of ten *Mesoplasma* strains using pDM023. * = below detection limit. **C)** Transformation efficiency of *M. florum* L1 compared to *M. florum* W20 as well as *M. entomophilum* W17. A significant difference was observed between L1 and W20 (p < 0.01).

Transformation efficiency varied by more than three orders of magnitude across the strains tested (Fig. 4B; Table S1). In the initial screen, *M. florum* W20, *M. entomophilum* W17, and *M. florum* L1 produced the highest transformation efficiencies, yielding 2.8 × 10², 2.3 × 10², and 1.5 × 10² transformants µg⁻¹ DNA, respectively. Several other strains produced few or no detectable transformants under the same conditions. These results demonstrate substantial variation in transformation efficiency even among closely related members of the *Mesoplasma* lineage.

We next examined *M. florum* W20 and *M. entomophilum* W17 under additional culture conditions, including cultures harvested at different growth stages, as inferred from medium coloration. W20 grew reproducibly at 34 °C and remained transformable, although variation in the selected growth phase did not produce a clear improvement in transformation efficiency relative to the initial screen (Fig. 4C; Table S1).

Initial cultures of W17 showed inconsistent growth under the conditions used for *M. florum* L1. Incubation at 30 °C restored reliable propagation, and transformation assays were therefore repeated at this temperature using cultures harvested at different growth stages. Under these adjusted conditions, W17 produced nearly two orders of magnitude more transformants than in the initial experiments (Fig. 4C; Table S1). Its transformation efficiency did not differ significantly from that of *M. florum* L1 (Welch’s two-sample *t*-test, *p* = 0.38). The improved performance of W17 following adjustment of its cultivation conditions further demonstrates the importance of recipient physiology in determining transformation efficiency.

The favorable transformation performance of W17, together with its closer phylogenetic relationship to *M. florum* than that of *M. capricolum*, identified this strain as a promising candidate for further characterization as a genome-transplantation recipient. We therefore examined whether W17 could support autonomous replication of plasmids carrying the *M. florum oriC*.

### *M. entomophilum* W17 supports replication of plasmids carrying the *M. florum oriC*

Having established suitable cultivation and transformation conditions for *M. entomophilum* W17, we next examined whether this strain could support autonomous replication of plasmids carrying chromosomal origins from different Mollicutes. We used a panel of plasmids containing the *oriC* and adjacent *dnaA* region from species previously examined in genome-transplantation experiments using *M. capricolum* as the recipient and in transformation into *M. florum* L1.^8^ The panel included two constructs sharing the same plasmid backbone but carrying the replication regions from *M. florum* L1 (pMflT-o4) or *M. entomophilum* W17 (pMenT-o4), as well as constructs carrying replication regions from *M. capricolum* subsp. *capricolum* (pMcapT), *M. mycoides* subsp. *mycoides* (pMmmT), *M. mycoides* subsp. *capricolum* (pMmcT), and two variants from *Spiroplasma citri* (pSciT-o2 and pSciT-o4) (Fig. S4). Each plasmid was transformed into *M. florum* L1 and *M. entomophilum* W17, allowing the compatibility of the different replication regions to be compared between the two species.

Only pMflT-o4 and pMenT-o4 produced detectable transformants in either host. In *M. florum* L1, pMflT-o4 transformed efficiently, whereas pMenT-o4 remained below the detection limit (Fig. 5A). None of the other heterologous *oriC* constructs produced detectable transformants. These results indicate that the plasmid carrying the native *M. florum oriC* was maintained in L1, whereas the replication regions from W17 and the more distantly related Mollicutes did not support detectable plasmid maintenance under the conditions tested.

**Figure 5:**
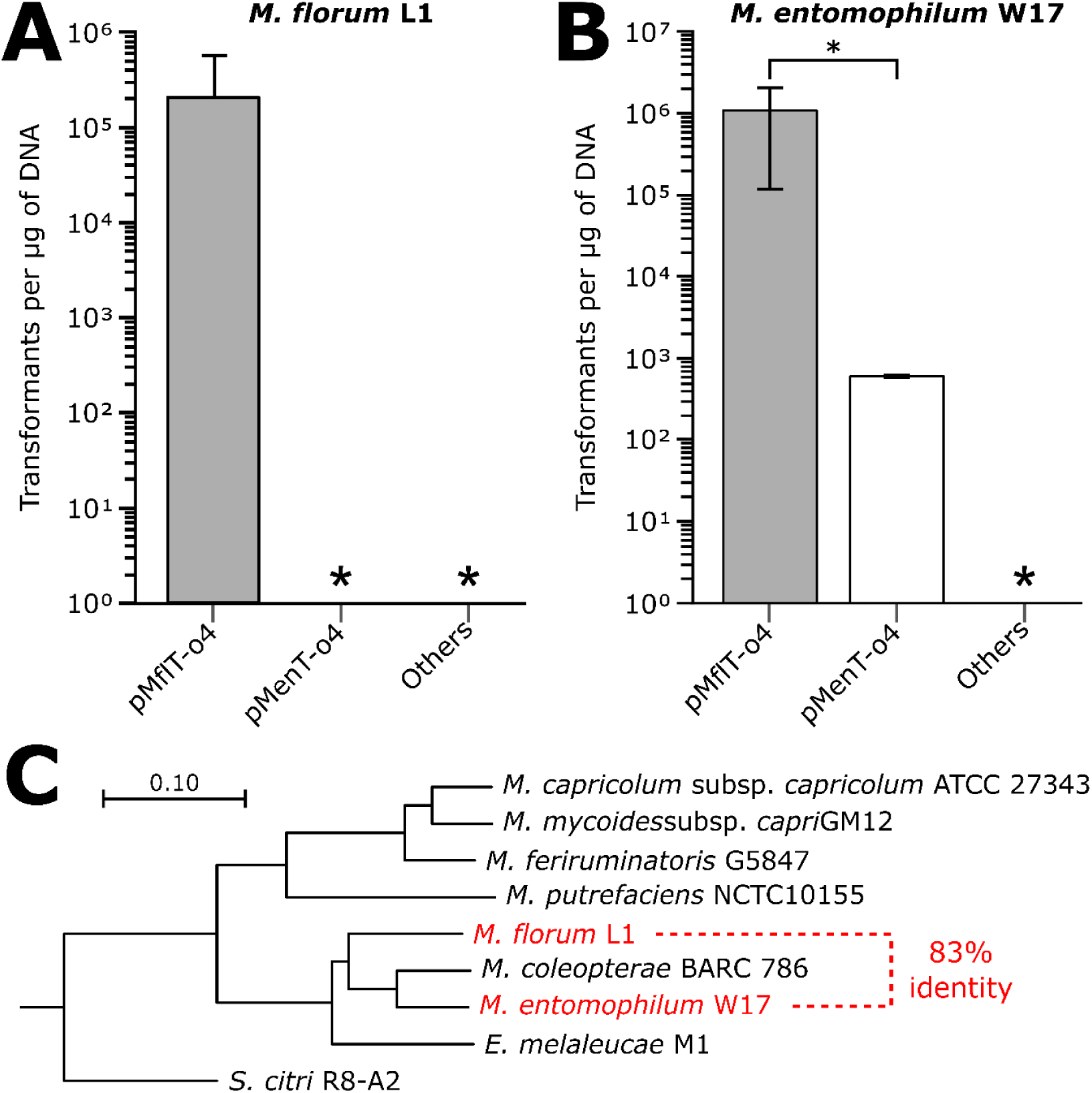
Compatibility of heterologous *oriC* plasmids in *M. florum* L1 and *M. entomophilum* W17. Transformation efficiency of plasmids carrying *oriC* regions from *M. florum*, *M. entomophilum* W17 or other Mollicutes species *M. capricolum* subsp. *capricolum*, *M. mycoides* subsp. *mycoides*, *M. mycoides* subsp. *capri* and *S. citri*) in *M. florum* L1 (A) and *M. entomophilum* W17 (B). * = below detection limit. Data in *M. florum* L1 were taken from Matteau *et al.*^12^, with the exception of pMenT- o4. **C)** Phylogenetic tree of the same *oriC* regions, aligned with MUSCLE 3.8.31 and inferred by maximum likelihood with PhyML 3.1 (see Methods).

By contrast, both pMflT-o4 and pMenT-o4 produced transformants in *M. entomophilum* W17 (Fig. 5B). Unexpectedly, pMflT-o4 yielded a higher transformation efficiency than pMenT-o4, even though the latter carries the native *M. entomophilum* W17 *oriC* and adjacent *dnaA* region. This difference was statistically significant (Welch’s two-sample *t*-test, *p* = 0.033). *M. entomophilum* W17 therefore displayed asymmetric compatibility with these replication regions: it supported plasmids carrying either the *M. entomophilum* W17 or *M. florum oriC*, whereas *M. florum* L1 supported only the construct carrying its native origin.

The *oriC* regions of *M. florum* L1 and *M. entomophilum* W17 share 83% sequence identity, which may contribute to the ability of pMflT-o4 to replicate in *M. entomophilum* W17 (Fig. 5C). This sequence similarity does not, however, explain why pMflT-o4 produced more transformants than the construct carrying the native *M. entomophilum* W17 replication region. Differences in the boundaries or organization of the cloned regions, the arrangement of DnaA-binding sites, the topology of DNA preparations or other plasmid features may influence their relative performance. Regardless of the underlying explanation, the ability of *M. entomophilum* W17 to support autonomous replication of a plasmid carrying the *M. florum oriC* demonstrates functional compatibility between the replication systems of the two species.

Together with its favorable transformation performance and closer phylogenetic relationship to *M. florum*, this replication compatibility strengthens the case for *M. entomophilum* W17 as a candidate recipient for transplantation of the *M. florum* L1 genome. These plasmid-level results provide a clear rationale for future experiments testing whether *M. entomophilum* W17 can receive and boot an intact *M. florum* chromosome.

### Implications for future genome transplantation

The transformation performance and replication compatibility of *M. entomophilum* W17 support its further evaluation as a recipient for transplantation of the *M. florum* L1 genome. Transformation efficiency alone is unlikely to determine recipient suitability, but the ability of *M. entomophilum* W17 to maintain plasmids carrying the *M. florum oriC* indicates functional compatibility between the replication systems of the two species. Using a heterologous recipient also reduces the extent of sequence homology between the donor and recipient chromosomes, thereby limiting homologous recombination that could generate chimeric genomes rather than cells controlled by the intact donor chromosome. An effective recipient may therefore need to combine sufficient physiological and molecular compatibility with enough sequence divergence to minimize recombination between the donor and recipient genomes. *M. entomophilum* W17 may offer such a balance because it is more closely related to *M. florum* than *M. capricolum* while remaining a distinct species.

A future transplantation workflow could involve the assembly and amplification of natural or synthetic *M. florum* chromosomes, followed by their transfer into *M. entomophilum* W17 and selection for cells controlled by the donor genome (Fig. 6). Synthetic chromosomes could include reduced, minimal, or recoded genome variants designed to investigate gene essentiality, genome organization, and the fundamental requirements of cellular life. Increasing transplantation efficiency beyond the few events currently obtained would facilitate the evaluation of multiple genome designs and enable more systematic or combinatorial approaches to synthetic genome construction.

**Figure 6:**
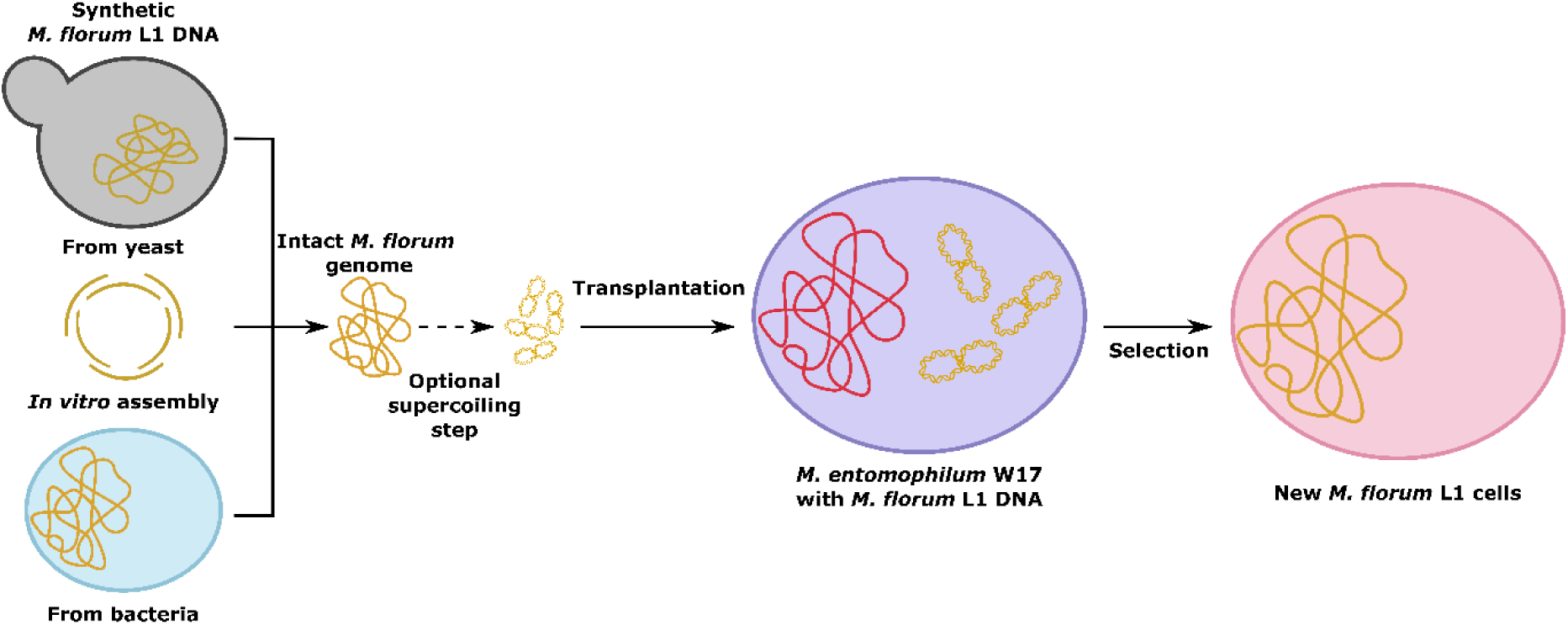
Conceptual model for generating synthetic *M. florum* cells using *M. entomophilum* W17. Synthetic *M. florum* L1 DNA is produced and compacted into a supercoiled form, transplanted into *M. entomophilum* W17, and selected to yield new cells derived from the synthetic genome.,

The pronounced effect of OriCiro-amplified plasmid DNA also suggests that the physical state of donor DNA should be considered when developing whole-genome transplantation protocols. Intact bacterial chromosomes face substantially greater physical constraints than plasmids because of their size and susceptibility to damage during preparation and transfer. Methods that preserve chromosome integrity and promote a compact DNA conformation may therefore improve the probability of successful delivery. Although the commercial OriCiro system is no longer available, the group that developed it has since described a cell-free transcription–translation (TXTL) approach in which a single plasmid encodes the proteins required to reconstitute *E. coli oriC*-dependent replication. This approach could provide an alternative route for generating replicated and highly supercoiled donor DNA without relying on the discontinued commercial kit.^30^ However, the effect observed with OriCiro-amplified pJG001 cannot be assumed to extend directly to complete bacterial chromosomes and will need to be evaluated using genome-sized donor DNA.

Recipient physiology represents a second important consideration. The strong effects of growth phase, cultivation temperature, membrane conditioning, and PEG concentration observed in this study indicate that transplantation conditions will need to be optimized specifically for *M. entomophilum* W17 rather than transferred directly from protocols developed for *M. florum* L1 or *M. capricolum*. In particular, the improved growth and transformation of *M. entomophilum* W17 at 30 °C illustrate how species-specific cultivation conditions can determine whether a strain displays favorable transformation properties.

This proposed framework positions *M. entomophilum* W17 as a promising recipient for future transplantation of the *M. florum* genome. Successful transplantation will require an intact *M. florum* chromosome to enter *M. entomophilum* W17, initiate replication, direct expression of the donor genome, and ultimately replace or eliminate the resident chromosome. Nevertheless, the combination of favorable transformation performance, phylogenetic proximity, functional compatibility with the *M. florum oriC*, and sufficient sequence divergence to reduce homologous recombination identifies *M. entomophilum* W17 as the strongest candidate emerging from this study for future genome-transplantation experiments.

## Methods

### Strains and culture conditions

All strains used in this study are listed in Table S2. *Mesoplasma* strains were cultured in ATCC 1161 medium at 34°C with shaking at 180-250 rpm, unless otherwise specified. ATCC 1161 medium was prepared as described previously.^8^ For *Mesoplasma* species, antibiotics were used at the following concentrations: streptomycin, 100 μg/ml; spectinomycin, 100 μg/ml; and tetracycline, 15 µg/ml. *E. coli* was grown in Luria-Bertani (LB) at 37°C with shaking at 180-250 rpm. Antibiotics were used at the following concentrations: ampicillin, 100 μg/ml; chloramphenicol, 200 μg/ml; streptomycin, 50 μg/ml; spectinomycin, 100 μg/ml; and tetracycline, 15 µg/ml. Strain MFSC001 was generated previously^20^ by Tn4001-mediated delivery of a Bxb1 *attB1* landing site into *M. florum* L1.

### Plasmid construction

All plasmids used in this study are listed in Table S2. Complete annotated sequences are available at http://bioinfo.ccs.usherbrooke.ca/PEG_transformation_florum.html. pJG001 was constructed by inserting the *E. coli* chromosomal origin of replication (*oriC*), amplified by PCR from the OriCiro system, into pCC027_v3^20^ by standard restriction ligation cloning using T4 DNA ligase. The resulting assembly was transformed into *E. coli* DH5α and selected on LB agar with corresponding antibiotics. Plasmids pDM017 to pDM023, which differ in promoter and resistance- marker combinations, were assembled by Gibson assembly from PCR products and transformed into DH5α *pir+*. pMenT-o4 and pSciT-o2 were also assembled by Gibson assembly using PCR products, and transformed into *E. coli* EC100D *pir+*. Other plasmids carrying *oriC* regions from other Mollicutes were described previously.^8^ All plasmid constructions were transformed into chemically competent calcium chloride treated cells by the traditional heat-shock method. Plasmid sequences were verified by restriction digest analysis and confirmed by Sanger or Oxford nanopore sequencing. Integration of pJG001 in *M. florum* MFSC001 was confirmed by PCR screening (Fig. S1) and Oxford nanopore sequencing.

### PEG-mediated transformation of Mollicutes species

All transformation solutions were sterilized before use by filtration through a 0.2 µm membrane. The PEG-mediated transformation procedure used in this study was based on the procedure described previously for *M. florum* L1^8^, with modifications and tested parameters listed in Table S1. Briefly, for each transformation, either 200 µl, 1.0 mL or 10 mL of a liquid batch culture harvested at various growth phase points was transferred into a 1.5 mL microcentrifuge tube (Axygen MCT-175-C or VWR 87003-294) and centrifuged at 21,300 × *g* for 2 min at 10 °C. To transform various *Mesoplasma* strains in the same experiment, cultures grown in a 96-well plate was instead used. The cell pellet was washed once or twice with either 1 mL of ice-cold S/T buffer (10 mM Tris-HCl pH 6.5, 250 mM NaCl, supplemented or not with 10 mM EDTA), electroporation buffer (272 mM sucrose, 1 mM HEPES, pH 7.4), PBS1X, TFBI (100 mM RbCl, 50 mM MnCl_2_, 30 mM potassium acetate, 10 mM CaCl_2_, 15% glycerol, pH 5.8), or 0.1 M CaCl_2_ by pipetting thoroughly to disrupt cell aggregates. The washed pellet was then resuspended in a variable volume (typically 40 µL) of CaCl_2_, varying in concentration from 5 mM to 500 mM. Cells were incubated on ice for either 30 or 60 min. After CaCl_2_ treatment, a varying amount of DNA (usually between 1 and 10 µg, along with or without 1 µL of yeast tRNAs) was added directly to the cell suspension, incubated on ice for 0, 5 or 15 min, followed by the addition of 120 µl (or otherwise specified) volume of 2× MFB buffer (between 10% and 30% PEG 8000 or 6000 - USB 19959, USB 19966 or Sigma 81255, 20 mM Tris-HCl pH 6.5 and 250 mM NaCl). Plasmid DNA was either amplified using the OriCiro kit (OriCiro Genomics) or purified using the EZ-10 Spin Column Plasmid DNA Miniprep Kit (Bio Basic), the QIAGEN Plasmid Maxi Kit (Qiagen) or a home-made protocol consisting of a classic chloroform purification followed by a precipitation with isopropanol and ethanol (see Supporting information). When specified, plasmid DNA was treated with enzymes prior to transformation (see Molecular Biology methods). Following MFB buffer addition, tubes were mixed gently by tapping, and 80 µl of ATCC 1161 medium lacking horse serum (replaced by 0.4% NaCl) was added. The volume of ATCC 1161 medium lacking horse serum was adjusted according to the volume of 2× MFB buffer used, and the order of reagent incorporation was sometimes modified (see Table S1 for more details). The mixture was incubated for 50 min at 34 °C (or 30°C) without agitation. Cells were then recovered by adding 1 ml of pre-warmed ATCC 1161 medium and incubating for 1h, 2 h or 3h at 34 °C without shaking. Following recovery, cells were centrifuged at maximum speed for 2 min at 10 °C and resuspended in the desired volume of ATCC 1161 medium. The suspension was plated on ATCC 1161 agar plates containing 200 U/mL penicillin and the appropriate selective antibiotics, and incubated at 34°C for 2 to 4 days. In parallel, serial dilutions from 10⁻¹ to 10⁻⁷ were spotted on the same medium without selective antibiotics to determine final cell concentrations. Other protocol modifications not listed above can be found in Table S1. The complete optimized protocol for *M. florum* L1 is available on protocols.io https://www.protocols.io/private/903EA367B76711F19CE90A58A9FEAC02.

### Molecular biology methods

PCR amplifications were performed using primers synthesized by either IDT or Twist Bioscience. Reactions were carried out with several DNA polymerases routinely used in the laboratory, mostly VeraSeq 2.0 High-Fidelity Polymerase (Qiagen), TransStart FastPfu Fly DNA Polymerase (TransGen), Taq-B DNA Polymerase (Qiagen), and OneTaq DNA Polymerase (NEB), each operated according to the manufacturer’s recommended conditions. Amplification products were assessed by electrophoresis on 0.7-1.2% agarose gels pre-stained with GelRed (Biotium) or Eco- Stain (Bio Basic) added at recommended concentrations. PCR amplifications were purified using either Agencourt AMPure XP beads (Beckman-Coulter) or the Zymo DNA Clean & Concentrator- 5 (Zymo Research). gDNA extraction was performed using the Quick-DNA Miniprep Kit (Zymo Research) following manufacturer’s specifications, or using Chelex 100 Resin on liquid cultures (Bio-Rad). Plasmid DNA purification was performed using either the EZ-10 Spin Column Plasmid DNA Miniprep Kit (Bio Basic), the QIAGEN Plasmid Maxi Kit (Qiagen) or a home-made protocol consisting of a classic chloroform purification followed by a precipitation with isopropanol and ethanol (see Supporting information). DNA preparations were either left untreated or subjected to one of the following treatments: topoisomerase I (NEB), gyrase (NEB), T4 DNA ligase (NEB) followed by gyrase, or linearization with a restriction enzyme followed by religation with T4 DNA ligase (NEB). DNA quantification was carried out using either a Nanodrop 2000 (Thermo Fisher) or the Quant-iT PicoGreen assay kit (Thermo Fisher). Gibson assembly reactions were performed using NEBuilder HiFi DNA Assembly Master Mix (NEB) according to the manufacturer’s specifications. OriCiro components were handled following the supplier’s protocol (OriCiro Genomics) and purified using Agencourt AMPure XP beads (Beckman-Coulter).

### Statistical analysis

Statistical significance was assessed using either one-way ANOVA or two-sample t-tests assuming unequal variances (Welch’s t-test), depending on the comparison. For each analysis, only the resulting p-value was used to determine significance, with a threshold of α = 0.05. All p-values reported in the Results section correspond directly to these tests and were calculated from untransformed transformation efficiency data (see Supporting information).

### Phylogenetic tree construction

*oriC* regions of selected Mollicute species were aligned with MUSCLE 3.8.31 and the tree inferred by maximum likelihood using PhyML 3.1, as described previously.^8^

## Abbreviations

PEG: polyethylene glycol

*oriC*: origin of chromosomal replication

## Author Information

Corresponding Author: Sébastien Rodrigue − Département de biologie, Université de Sherbrooke, Sherbrooke, Québec J1K 2R1, Canada; orcid.org/0000-0002-5366- 7234;

## Authors

Jérémy Gagnon - Département de biologie, Université de Sherbrooke, Sherbrooke, Québec J1K 2R1, Canada; orcid.org/0009-0009-3460-8376

Dominick Matteau - Département de biologie, Université de Sherbrooke, Sherbrooke, Québec J1K 2R1, Canada; orcid.org/0009-0005-2975-4687

Frederic Grenier - Département de biologie, Université de Sherbrooke, Sherbrooke, Québec J1K 2R1, Canada; orcid.org/0000-0003-3910-3165

Sébastien Rodrigue - Département de biologie, Université de Sherbrooke, Sherbrooke, Québec J1K 2R1, Canada; orcid.org/0000-0002-5366-7234

JG: Conceptualization, Data curation, Formal analysis, Investigation, Methodology, Validation, Visualization, Writing - original draft, Writing - review and editing

DM: Conceptualization, Data curation, Formal analysis, Investigation, Methodology, Supervision, Validation, Visualization, Writing - review and editing FG: Supervision, Writing - review and editing

SR: Project administration, Conceptualization, Supervision, Resources, Funding Acquisition, Writing - review and editing

## Acknowledgment

We thank Catherine Chamberland for providing the *M. florum* MFSC001 strain and the members of the Rodrigue laboratory for technical assistance, sequencing, and helpful discussions throughout this work. This work was supported by the Natural Sciences and Engineering Research Council of Canada (NSERC).

---

**Figure S1:**
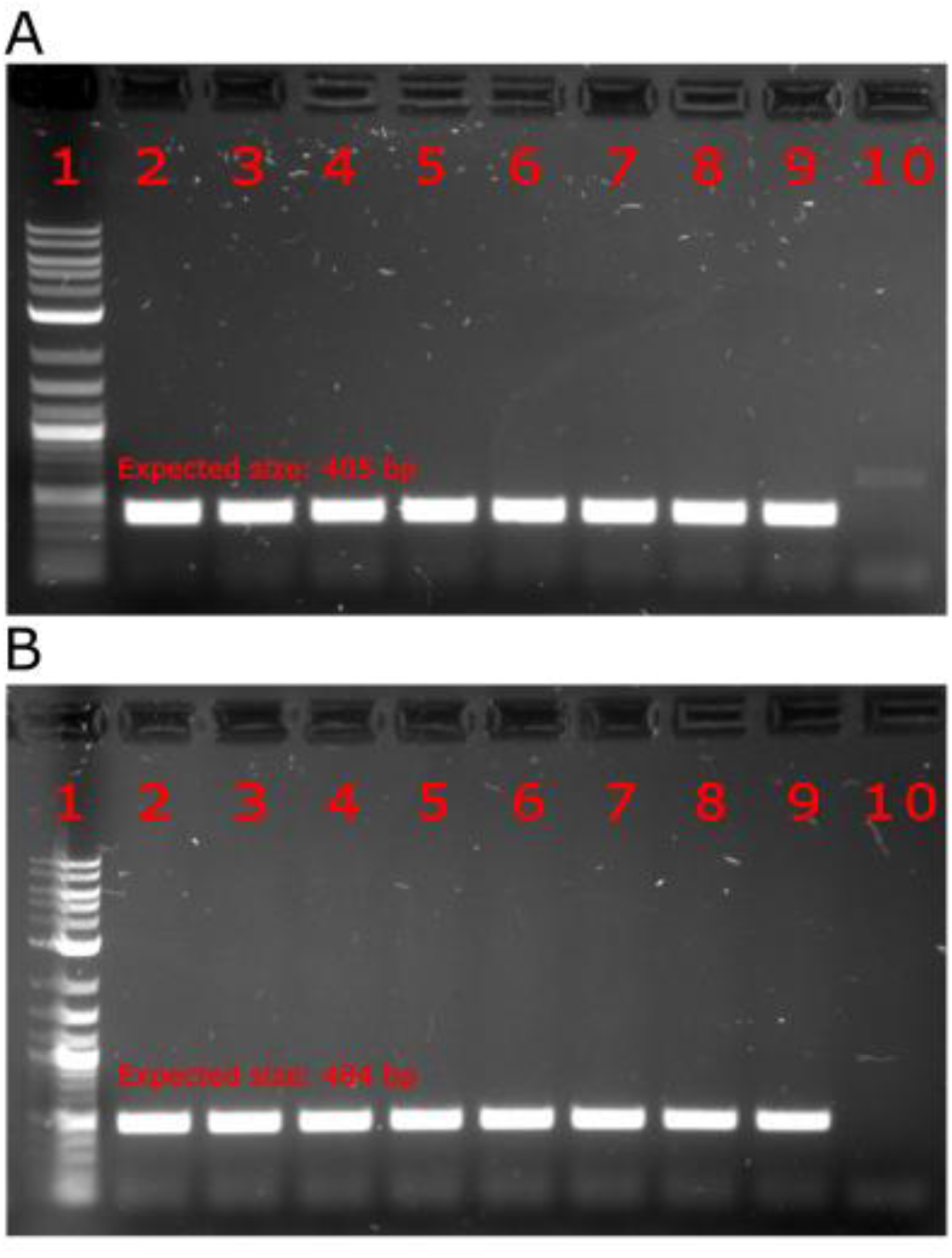
PCR confirmation of pJG001 integration in *M. florum* MFSC001. PCR amplification of the left (A) and right (B) integration borders across 8 different clones (lanes 2-9). Lanes 1 and 10 contain the 1 kb Plus DNA Ladder (NEB N3200) and a negative control (*M. florum* MFSC001), respectively. Expected size: 405 bp (A) and 484 bp (B) if integrated.

**Figure S2.**
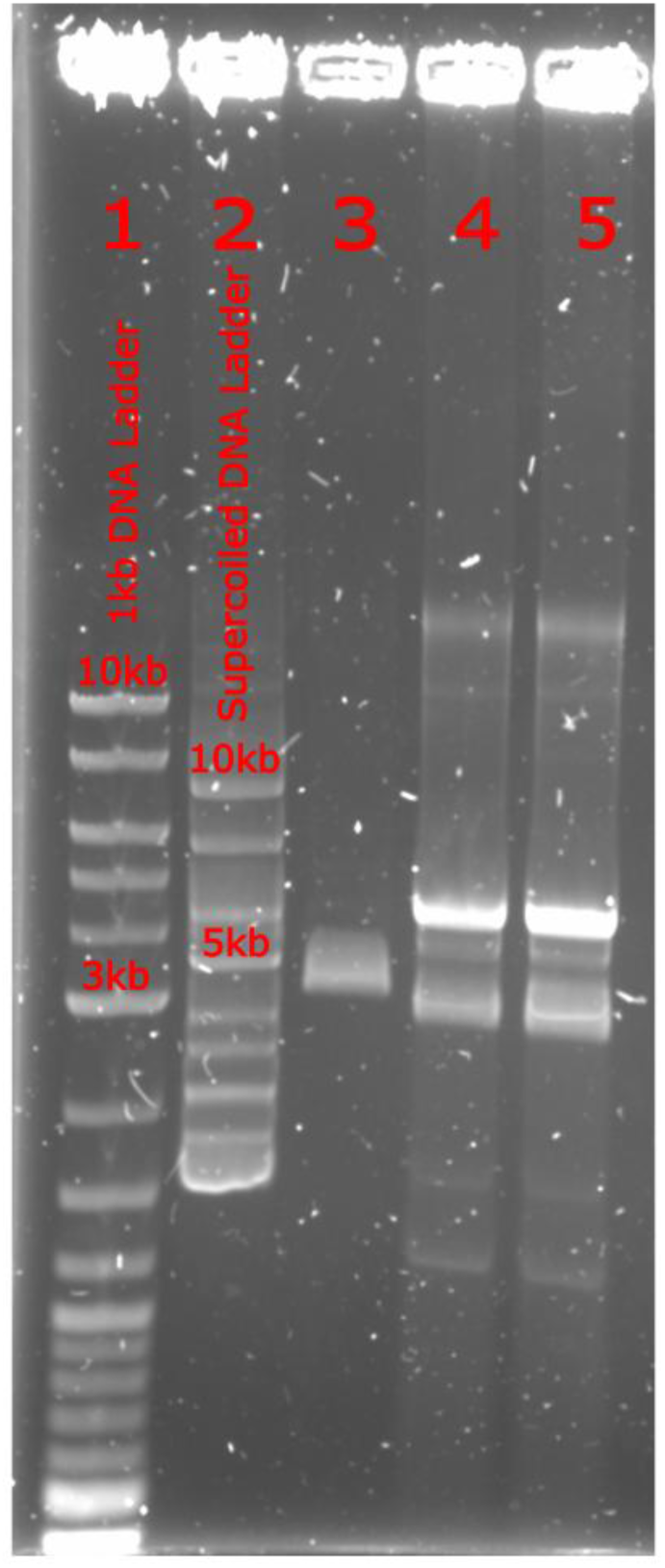
OriCiro treatment compared to topoisomerase I and Maxiprep conditions. Electrophoresis results of pJG001 plasmid amplified using OriCiro or treated with topoisomerase I migrated on a 0.7% agarose gel for 75min at 130V. Lane 1: 1 kb Plus DNA Ladder (NEB N3200); lane 2: Supercoiled DNA Ladder (NEB N0472); lane 3: OriCiro pJG001 SPRI-purified; lane 4: Maxiprep pJG001, topoisomerase I treated and SPRI-purified; lane 5: Maxiprep pJG001 SPRI-purified. While maxiprep-prepared plasmid DNA yielded multiple bands on gel, indicating various DNA topologies (nicked, linear, supercoiled), OriCiro pJG001 showed a single band corroborating previous observations (citations).

**Figure S3.**
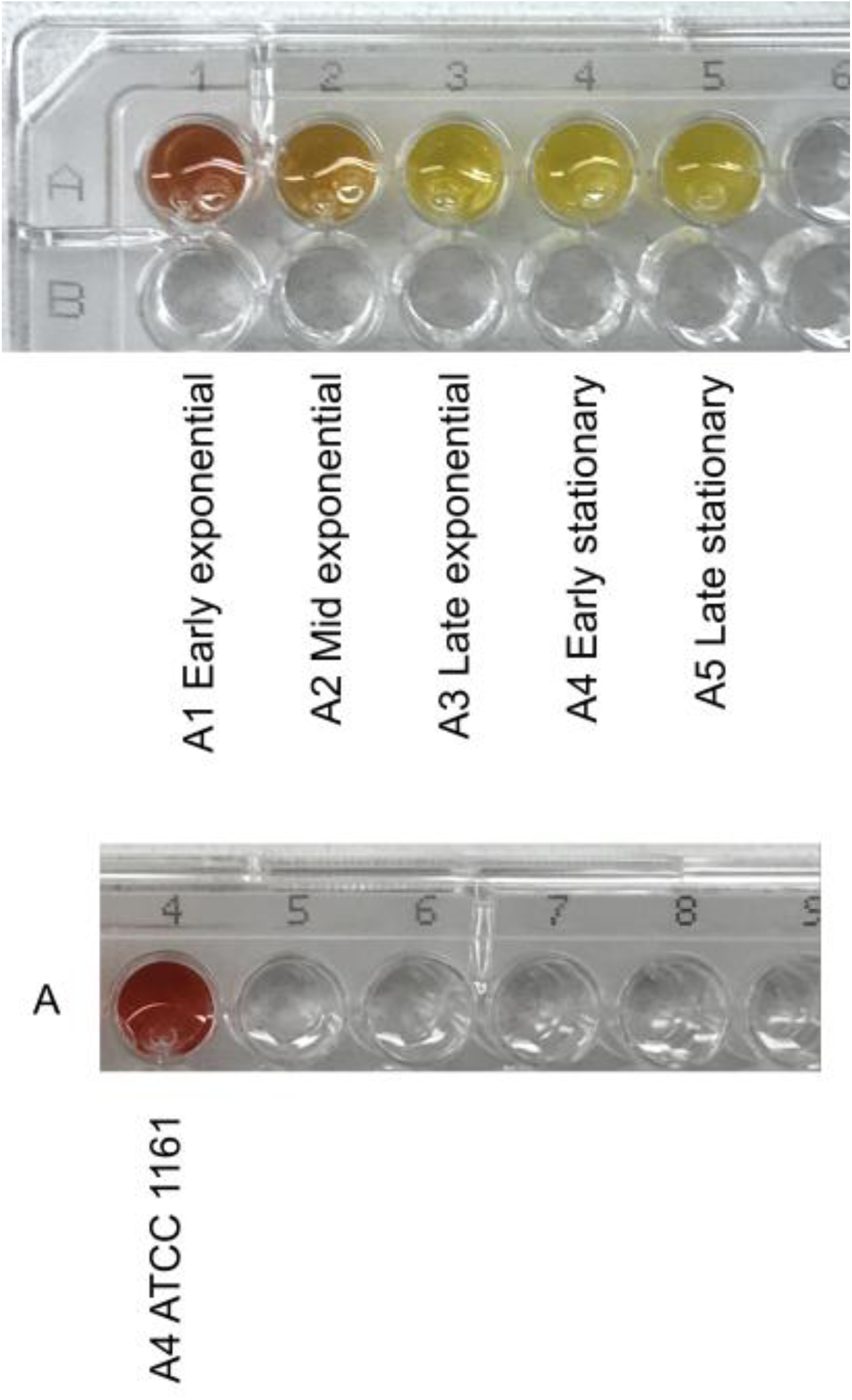
**Culture appearance across *M. florum* growth stages**. Cultures at five stages of growth, defined by phenol red coloration: early exponential (dark orange, A1), mid exponential (pale orange, A2), late exponential (translucent yellow, A3), early stationary (cloudy yellow, A4), and late stationary (opaque yellow, A5). Bottom panel: non inoculated ATCC 1161 media, A4.

**Figure S4:**
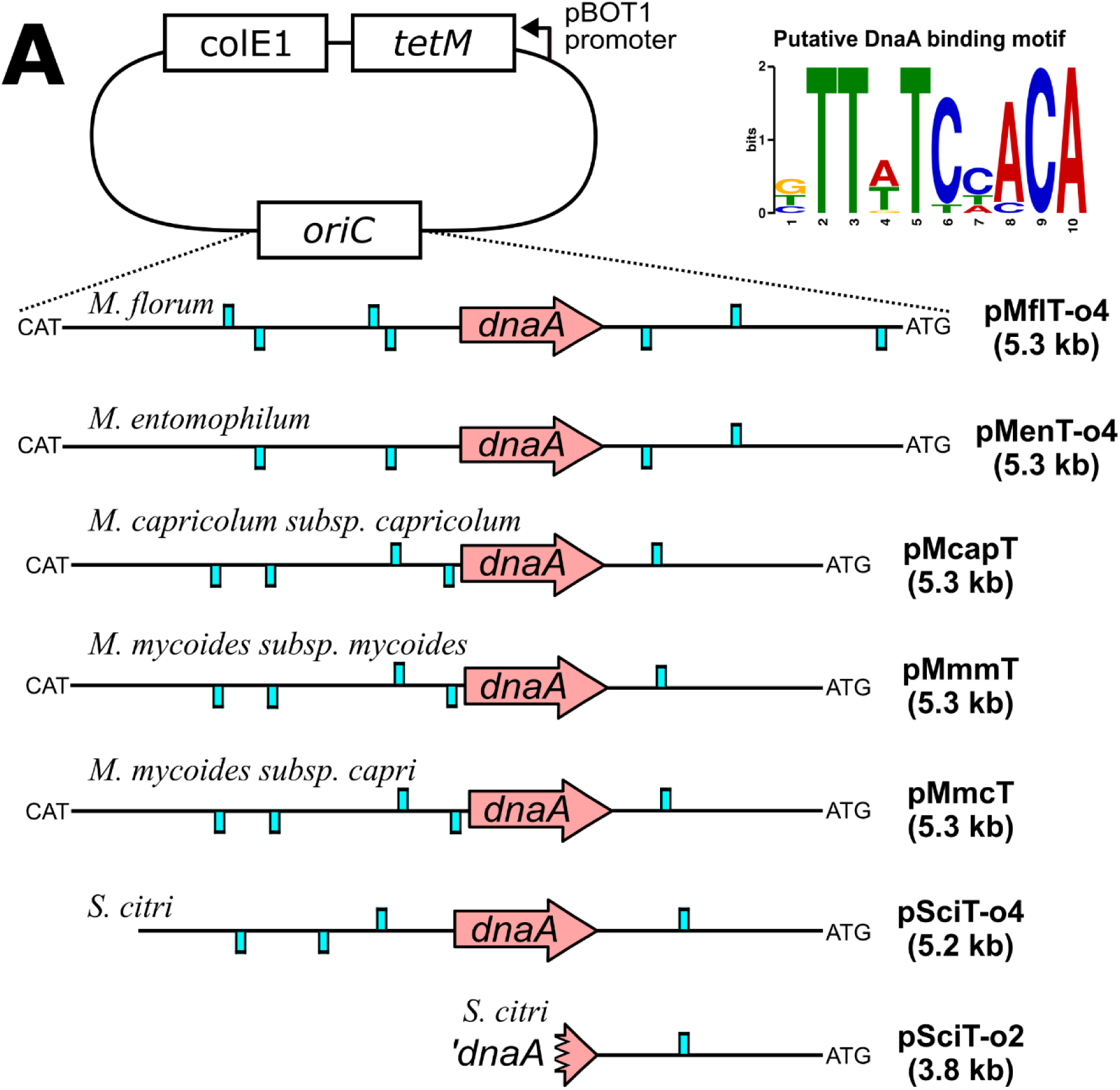
Plasmids carrying heterologous *oriC* regions. Schematic representation of *oriC* plasmids described previously (adapted from Matteau *et al*.^8^), together with pMflT-o4 (*M. florum* L1), pMenT-o4 (*M. entomophilum* W17) and pSciT-o2 (*Spiroplasma citri*). Pale blue boxes indicate putative DnaA binding sites across each *oriC* region.

**Table S1:** Source data for PEG-mediated transformation experiments.

**Table S2:** Strains and plasmids used in this study.

