## Supporting information for "Optimization of DNA Transformation in *Mesoplasma florum* and Identification of a Candidate Recipient Strain for Genome Transplantation"

**Statistical analyses :**

Figure 3B statistical analysis

| t-Test: Two-Sample Assuming Unequal Variances |  |  |
| --- | --- | --- |
|  | *Variable 1* | *Variable 2* |
| Mean | 25052.5 | 120 |
| Variance | 3939624.5 | 36266.66667 |
| Observations | 2 | 4 |
| Hypothesized Mean Difference | 0 |  |
| df | 1 |  |
| t Stat | 17.72377455 |  |
| P(T<=t) one-tail | 0.01794046537 |  |
| t Critical one-tail | 6.313751515 |  |
| P(T<=t) two-tail | 0.03588093074 |  |
| t Critical two-tail | 12.70620474 |  |

Figure 3C statistical analysis

| Anova: Single Factor |  |  |  |  |  |  |
| --- | --- | --- | --- | --- | --- | --- |
| SUMMARY |  |  |  |  |  |  |
| *Groups* | *Count* | *Sum* | *Average* | *Variance* |  |  |
| Ligase + Gyrase | 3 | 9844 | 3281.333333 | 3553640.333 |  |  |
| Gyrase | 3 | 8711 | 2903.666667 | 2599334.333 |  |  |
| Linearisation + Ligation | 3 | 5433 | 1811 | 811447 |  |  |
| Topoisomerase | 3 | 8745 | 2915 | 2599543 |  |  |
| Maxiprep | 3 | 10901 | 3633.666667 | 1151396.333 |  |  |
| ANOVA |  |  |  |  |  |  |
| *Source of Variation* | *SS* | *df* | *MS* | *F* | *P-value* | *F crit* |
| Between Groups | 5608326.933 | 4 | 1402081.733 | 0.6542391494 | 0.63713999 | 3.478049691 |
| Within Groups | 21430722 | 10 | 2143072.2 |  |  |  |
| Total | 27039048.93 | 14 |  |  |  |  |

Figure 3D statistical analysis

| Anova: Single Factor |  |  |  |  |  |  |
| --- | --- | --- | --- | --- | --- | --- |
| SUMMARY |  |  |  |  |  |  |
| *Groups* | *Count* | *Sum* | *Average* | *Variance* |  |  |
| Ligase + Gyrase | 3 | 64222 | 21407.33333 | 1311352897 |  |  |
| Gyrase | 3 | 134345 | 44781.66667 | 5647342476 |  |  |
| Topoisomerase | 2 | 74223 | 37111.5 | 2237069161 |  |  |
| Linearisation + Ligation | 2 | 26000 | 13000 | 332252642 |  |  |
| Maxiprep | 3 | 108412 | 36137.33333 | 3533723961 |  |  |
| ANOVA |  |  |  |  |  |  |
| *Source of Variation* | *SS* | *df* | *MS* | *F* | *P-value* | *F crit* |
| Between Groups | 1646380000 | 4 | 411595000 | 0.1397952605 | 0.9626267985 | 3.837853355 |
| Within Groups | 23554160473 | 8 | 2944270059 |  |  |  |
| Total | 25200540472 | 12 |  |  |  |  |

Figure 4B statistical analysis compared to L1

| t-Test: Two-Sample Assuming Unequal Variances |  |  |
| --- | --- | --- |
|  | *L1* | *W20* |
| Mean | 152.5 | 282.5 |
| Variance | 23112.5 | 15312.5 |
| Observations | 2 | 2 |
| Hypothesized Mean Difference | 0 |  |
| df | 2 |  |
| t Stat | -0.9378889347 |  |
| P(T<=t) one-tail | 0.2236543419 |  |
| t Critical one-tail | 2.91998558 |  |
| P(T<=t) two-tail | 0.4473086839 |  |
| t Critical two-tail | 4.30265273 |  |

| t-Test: Two-Sample Assuming Unequal Variances |  |  |
| --- | --- | --- |
|  | *L1* | *W17* |
| Mean | 152.5 | 227.5 |
| Variance | 23112.5 | 1012.5 |
| Observations | 2 | 2 |
| Hypothesized Mean Difference | 0 |  |
| df | 1 |  |
| t Stat | -0.6828771804 |  |
| P(T<=t) one-tail | 0.3092873487 |  |
| t Critical one-tail | 6.313751515 |  |
| P(T<=t) two-tail | 0.6185746974 |  |
| t Critical two-tail | 12.70620474 |  |

| t-Test: Two-Sample Assuming Unequal Variances |  |  |
| --- | --- | --- |
|  | *L1* | *W37* |
| Mean | 152.5 | 47.5 |
| Variance | 23112.5 | 4512.5 |
| Observations | 2 | 2 |
| Hypothesized Mean Difference | 0 |  |
| df | 1 |  |
| t Stat | 0.8934148226 |  |
| P(T<=t) one-tail | 0.2678995889 |  |
| t Critical one-tail | 6.313751515 |  |
| P(T<=t) two-tail | 0.5357991778 |  |
| t Critical two-tail | 12.70620474 |  |

| t-Test: Two-Sample Assuming Unequal Variances |  |  |
| --- | --- | --- |
|  | *L1* | *MouA-2* |
| Mean | 152.5 | 22.5 |
| Variance | 23112.5 | 1012.5 |
| Observations | 2 | 2 |
| Hypothesized Mean Difference | 0 |  |
| df | 1 |  |
| t Stat | 1.183653779 |  |
| P(T<=t) one-tail | 0.223291755 |  |
| t Critical one-tail | 6.313751515 |  |
| P(T<=t) two-tail | 0.4465835099 |  |
| t Critical two-tail | 12.70620474 |  |

| t-Test: Two-Sample Assuming Unequal Variances |  |  |
| --- | --- | --- |
|  | *L1* | *W23* |
| Mean | 152.5 | 10 |
| Variance | 23112.5 | 200 |
| Observations | 2 | 2 |
| Hypothesized Mean Difference | 0 |  |
| df | 1 |  |
| t Stat | 1.319883007 |  |
| P(T<=t) one-tail | 0.2063840617 |  |
| t Critical one-tail | 6.313751515 |  |
| P(T<=t) two-tail | 0.4127681235 |  |
| t Critical two-tail | 12.70620474 |  |

| t-Test: Two-Sample Assuming Unequal Variances |  |  |
| --- | --- | --- |
|  | *L1* | *BARC787* |
| Mean | 152.5 | 5 |
| Variance | 23112.5 | 50 |
| Observations | 2 | 2 |
| Hypothesized Mean Difference | 0 |  |
| df | 1 |  |
| t Stat | 1.370611281 |  |
| P(T<=t) one-tail | 0.200636362 |  |
| t Critical one-tail | 6.313751515 |  |
| P(T<=t) two-tail | 0.4012727241 |  |
| t Critical two-tail | 12.70620474 |  |

| t-Test: Two-Sample Assuming Unequal Variances |  |  |
| --- | --- | --- |
|  | *L1* | *BARC786* |
| Mean | 152.5 | 2.5 |
| Variance | 23112.5 | 12.5 |
| Observations | 2 | 2 |
| Hypothesized Mean Difference | 0 |  |
| df | 1 |  |
| t Stat | 1.394971665 |  |
| P(T<=t) one-tail | 0.1979735645 |  |
| t Critical one-tail | 6.313751515 |  |
| P(T<=t) two-tail | 0.3959471291 |  |
| t Critical two-tail | 12.70620474 |  |

| t-Test: Two-Sample Assuming Unequal Variances |  |  |
| --- | --- | --- |
|  | *L1* | *BARC781* |
| Mean | 152.5 | 0 |
| Variance | 23112.5 | 0 |
| Observations | 2 | 2 |
| Hypothesized Mean Difference | 0 |  |
| df | 1 |  |
| t Stat | 1.418604651 |  |
| P(T<=t) one-tail | 0.1954483298 |  |
| t Critical one-tail | 6.313751515 |  |
| P(T<=t) two-tail | 0.3908966596 |  |
| t Critical two-tail | 12.70620474 |  |

| t-Test: Two-Sample Assuming Unequal Variances |  |  |
| --- | --- | --- |
|  | *L1* | *CnuA-2* |
| Mean | 152.5 | 0 |
| Variance | 23112.5 | 0 |
| Observations | 2 | 2 |
| Hypothesized Mean Difference | 0 |  |
| df | 1 |  |
| t Stat | 1.418604651 |  |
| P(T<=t) one-tail | 0.1954483298 |  |
| t Critical one-tail | 6.313751515 |  |
| P(T<=t) two-tail | 0.3908966596 |  |
| t Critical two-tail | 12.70620474 |  |

Figure 4C statistical analysis compared to L1

| t-Test: Two-Sample Assuming Unequal Variances |  |  |
| --- | --- | --- |
|  | *L1* | *W17* |
| Mean | 879.5416667 | 38148.33333 |
| Variance | 1079796.085 | 3252529082 |
| Observations | 24 | 3 |
| Hypothesized Mean Difference | 0 |  |
| df | 2 |  |
| t Stat | -1.131843378 |  |
| P(T<=t) one-tail | 0.1875729499 |  |
| t Critical one-tail | 2.91998558 |  |
| P(T<=t) two-tail | 0.3751458999 |  |
| t Critical two-tail | 4.30265273 |  |

| t-Test: Two-Sample Assuming Unequal Variances |  |  |
| --- | --- | --- |
|  | *L1* | *W20* |
| Mean | 879.5416667 | 169.3333333 |
| Variance | 1079796.085 | 19649.33333 |
| Observations | 24 | 3 |
| Hypothesized Mean Difference | 0 |  |
| df | 24 |  |
| t Stat | 3.128298298 |  |
| P(T<=t) one-tail | 0.002283133604 |  |
| t Critical one-tail | 1.71088208 |  |
| P(T<=t) two-tail | 0.004566267208 |  |
| t Critical two-tail | 2.063898562 |  |

Figure 5C statistical analysis

| t-Test: Two-Sample Assuming Unequal Variances |  |  |
| --- | --- | --- |
|  | *pMenT-04* | *pMflT-04* |
| Mean | 600 | 1066527.571 |
| Variance | 800 | 1051797544361 |
| Observations | 2 | 7 |
| Hypothesized Mean Difference | 0 |  |
| df | 6 |  |
| t Stat | -2.74986035 |  |
| P(T<=t) one-tail | 0.0166489443 |  |
| t Critical one-tail | 1.943180281 |  |
| P(T<=t) two-tail | 0.03329788861 |  |
| t Critical two-tail | 2.446911851 |  |

### **Supplementary Protocol: Home-made plasmid miniprep**

#### **Solutions**

**Solution I** (50 mM glucose, 25 mM Tris-HCl pH 8.0, 10 mM EDTA) — per 100 mL

- 0.9 g glucose
- 2.5 mL Tris-HCl 1 M, pH 8.0
- 2 mL EDTA 0.5 M
- Complete to 100 mL with distilled water. Store at 4 °C.

**Solution II** (1% SDS, 0.2 M NaOH) — per 100 mL

- 1 g SDS
- 20 mL NaOH 1 N
- Complete to 100 mL with distilled water. Sterilize by filtration and store at room temperature. Prepare fresh monthly.

**Solution III** (3 M potassium acetate) — per 100 mL

- 60 mL potassium acetate 5 M
- 11.5 mL glacial acetic acid
- 28.5 mL distilled water
- Store at 4°C.

**TE 1X** (10 mM Tris-HCl pH 8.0, 1 mM EDTA) — per 100 mL

- 1 mL Tris-HCl 1 M, pH 8.0
- 200 µL EDTA 0.5 M
- Complete to 100 mL with distilled water.

#### **Procedure**

1. Transfer 1.5 mL of overnight culture to a microtube and centrifuge at maximum speed. Remove the supernatant by aspiration.
2. Resuspend the pellet in 200 µL of Solution I and add 2 µL of RNase A (10 mg/ml).
3. Add 200 µL of Solution II and invert the tube five times. **Invert gently** — vigorous mixing shears genomic DNA and increases contamination of the final preparation.
4. Add 200 µL of Solution III and invert the tube five times. A white precipitate forms.
5. **In a fume hood**. add 600 µL of chloroform and vortex for a few seconds.
6. Centrifuge at maximum speed for 5 min. Handle the tube carefully afterwards to avoid disturbing the phases.
7. Transfer the upper aqueous phase to a clean microtube.
8. Add 600 µL of isopropanol and invert the tube ten times.
9. Centrifuge at maximum speed for 5 min and remove as much supernatant as possible.
10. Add 200 µL of 70% ethanol.
11. Centrifuge at maximum speed for 5 min, remove the supernatant, and air-dry the pellet at room temperature.
12. Resuspend the DNA in 50 µL of water or TE 1X.
